# Enhanced photo-crosslinking in living cells with high-intensity longwave ultraviolet light

**DOI:** 10.1101/2025.08.20.671286

**Authors:** Jakob Trendel, Polina Prokofeva, Zhuo Angel Chen, Lukas Horn, Simon Trendel, Mirea Mema, Marchel Stuiver, Juri Rappsilber, Bernhard Kuster

## Abstract

The activation of chemical reactions in living cells using ultraviolet (UV) light enables the interrogation of biomolecules in their native environment with photoreactive probes or crosslinking reagents. Although numerous photo-crosslinking approaches have been successfully employed, they often suffer from common limitations, including low reaction yields, the need for long exposure times, and irradiation-induced cellular damage from heat, desiccation, or side reactions. We recently showed that 365 nm light-emitting diodes (LEDs) enable rapid, bioorthogonal protein-DNA crosslinking in living cells, incurring minimal photodamage. Here we generalize this approach and demonstrate that high-intensity, longwave UV light reduces the irradiation time for in-cell photo-crosslinking reactions by up to 1000-fold, allowing protein-drug, protein-protein, protein-DNA and protein-RNA interactions to be fixed within seconds. Benchmarking this rapid photo-activation for the analysis of RNA-interacting proteomes responding to RNA-binding drugs, we demonstrate both qualitative and quantitative advantages of controlled, high-intensity UV irradiation, uncovering emergent experimental opportunities that were previously inaccessible to light-activated chemistry in intact cells and tissues.

## Introduction

The interaction of a biomolecule with a second biomolecule or a drug contains information that can indicate their relationship and function. For example, the interaction of a protein with DNA(1–4), RNA(5–11) or another protein(12, 13) can indicate the biological function of that protein as part of a complex or phase-separated compartment. Similarly, the binding of a kinase or HDAC inhibitor to a specific protein, can imply that the molecule inhibits the enzymatic function of this protein.(14, 15) Such an interaction occurs typically through combinations of many weak, non-covalent interactions between the participating molecules. To generate biochemical evidence for this interaction in living cells, photo-crosslinking can be applied (Figure 1A). To this end, a photoreactive group is introduced into one of the interaction partners, which, upon activation by light, creates an artificial, covalent bond between the two molecules. This photo-crosslinked complex can be purified from non-crosslinked, unspecific interactors using one of the complex partners as enrichment handle. The composition of the complex can subsequently be determined by proteomic, genomic or transcriptomic analysis.(1, 7–9, 16, 17) Because the linkage between the photo-crosslinked partners is covalent, purification of the complex can be strongly denaturing, leading to very effective elimination of unspecific or indirect interactors. Moreover, the covalent crosslinking site itself can lead to aberrations in the detection of protein or nucleic acids by mass spectrometry or sequencing, respectively, oftentimes pinpointing the exact interaction site in their sequence.(18–21)

**Figure 1:**
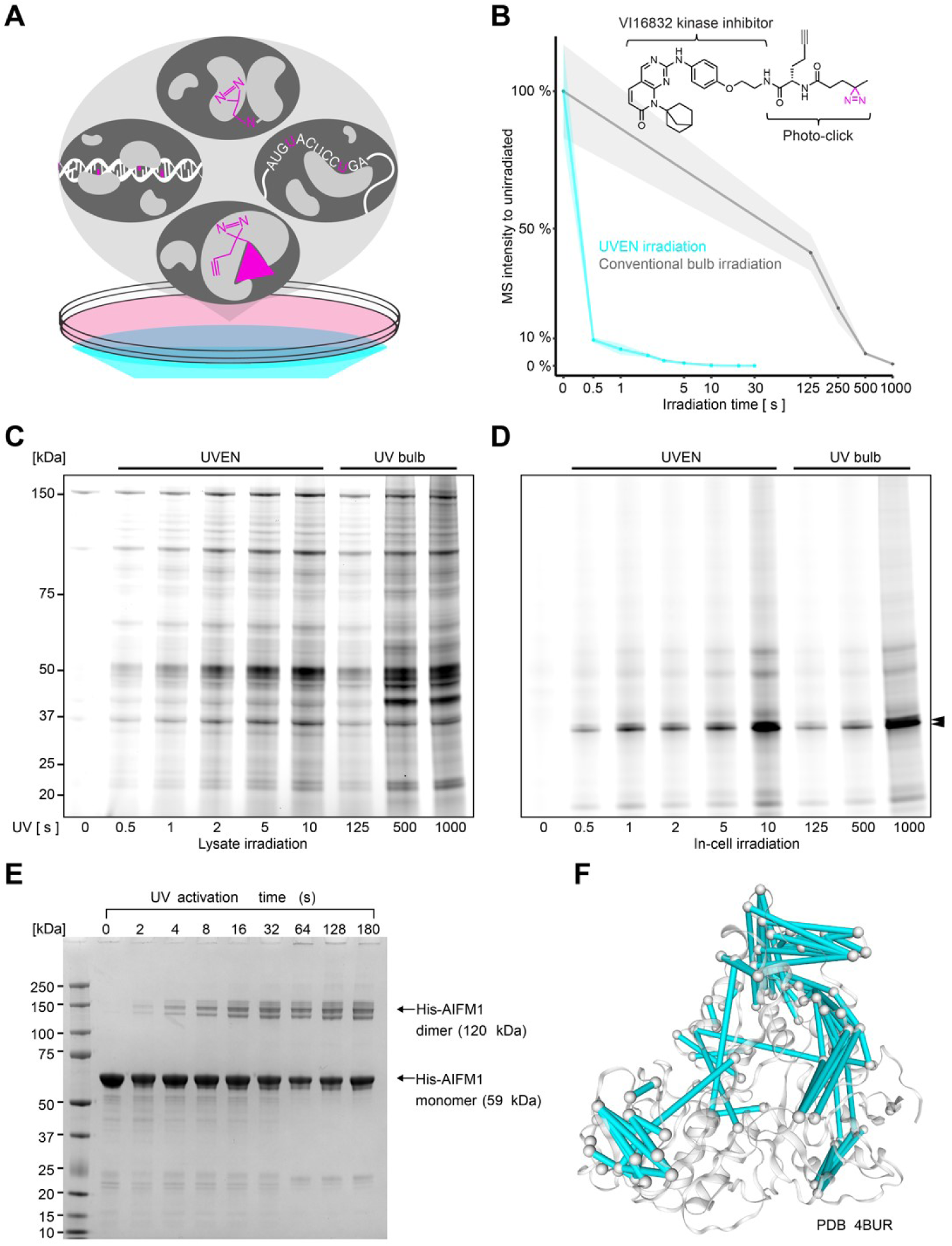
Rapid activation of crosslinking reactions using high-intensity UV irradiation. A) Schematic representation of four photo-crosslinking reactions presented in this study. Photoprobes can be metabolically incorporated into DNA, RNA, added ectopically in the form of photoreactive drugs or bifunctional protein-protein crosslinkers. Their activation occurs by irradiating culture cells with UV light. B) Line plot showing LC-MS quantification of a kinase inhibitor PAL probe upon irradiation in water. Compared are various irradiation times with a conventional bulb-based device (Vilber Biolink) or the LED-based UVEN. Irradiation occurred at identical distance from the irradiant (3.5 cm). C) Fluorescence imaging of SDS-PAGE after PAL in native lysates. Compared are various time points of LED irradiation (UVEN) to conventional bulb irradiation at identical distance. D) Same as in B but for PAL in living cells. Arrows indicate expected molecular weight of CDK4/6 (34/ 37 kDA). E) SDS-PAGE analysis of sulfo-SDA crosslinked AIFM1 with UV activation for 0-180 s by UVEN. A crosslinked dimeric AIFM1 band appears exclusively upon UV exposure. The yield of the dimeric crosslinked product increases with activation time and plateaus at 32 s. F) Visualization of sulfo-SDA crosslinks obtained after 32 s of UV activation, mapped onto the crystal structure of AIFM1 (PDB ID: 4BUR). A total of 86 crosslinks (87% of 99 displayable crosslinks) with Cα–Cα distances below 30 Å are shown as cyan lines connecting the Cα atoms of crosslinked residues.

Activation of a crosslinking reaction by light is conceptually powerful for cell biology because it is minimally invasive and can be timed. For example, for investigating protein-RNA interactions in culture cells, their transcriptome can be metabolically labeled with the photo-activatable nucleotide 4-thiouridine (4SU). Additional perturbations can be applied to investigate if proteins change their interaction with RNA between conditions and crosslinking triggered by 1-2 min of irradiation with UV bulbs placed directly above the cells. Thus, compared to the most commonly applied form of chemical crosslinking with formaldehyde, which requires 10 min of fixation at room temperature, photo-crosslinking is fast and carried out at 4 °C on ice, ‘freezing’ the state of the cell in the moment of crosslinking. Yet, effective protein crosslinking with other photoprobes, such as diazirine-modified small molecules, can require 10-20 min of irradiation or longer (22), and some photoprobes, such as the deoxyribonucleotide 4-thiothymidine, have such low reactivity that conventional bulb-based irradiation systems are often not sufficient.(17)

Here, we show that high-intensity UV irradiation from an LED-based device accelerates commonly used photoreactions in cells by several orders of magnitude, enabling us to perform protein-drug, protein-protein and protein-RNA crosslinking within seconds while maintaining cells in their original medium. Further assessing irradiation at physiological culture conditions, we demonstrate quantitative and qualitative advantages of high-intensity photo-activation, and evaluate rapid protein-RNA crosslinking on the RNA-interacting proteomes of breast cancer cells responding to a panel of RNA-binding drugs. In addition, we demonstrate protein-DNA crosslinking across different cell types and find temperature dependent crosslinking of C2H2-type zinc-finger proteins.

## Methods

### Cell culture

MCF7 (human female breast adenocarcinoma) and U2OS (human female bone sarcoma) were obtained from ATCC. U251 (human male glioblastoma) and HT-29 (human female colon carcinoma) were obtained as part of the NCI60 panel. Cells were maintained in Dulbecco’s Modified Eagle’s Medium (DMEM) supplemented with 10 % dialysed FBS and Pen-Strep (100 U / ml penicillin, 100 mg / ml streptomycin) at 37 °C, 5 % CO_2_. For the comparative analysis of DNA interactomes, 2 million cells were seeded and expanded in the presence of 100 µM 4-thiothymidine (4ST) over four days. For the comparative analysis of RNA interactomes, 2 million cells were expanded for three days and 100 µM 4-thiouridine (4SU) added four 24 h before UV crosslinking.

### Construction of the UVEN irradiation device

Housings of various prototypes were built from milled wood and 3D-printed plastic elements, which served as scaffold for the assembly of commercially available aluminum and electrical parts. High-intensity UV-LEDs (16 Luminous SBM-120-UV-F34-H365-22) were soldered on a custom copper-based PCB and mounted onto an aluminum heat sink. A publication on the construction of the device is currently in preparation. UVEN is presented as an open-science project, for which we provide detailed bills of materials, computer-aided designs, construction manuals as well as open-source firmware at www.uven.org.

### Operation of the UVEN irradiation device for UV crosslinking

For UV activation of photo-crosslinking reactions in cells cultured on 15 cm diameter culture dishes, the UVEN irradiation device was operated at 3.5 Ampere per LED resulting in 2,000 +/- 300 mW/cm^2^ intensity across the entire area of the culture dish. For protein-DNA crosslinking 30 s of UVEN irradiation was applied, where cells were washed with 50 ml ice-cold PBS and subsequently irradiated in another 50 ml PBS. For all other experiments with irradiation times up to 5 s, cells were placed from the incubator immediately into the irradiation device, remaining in their original medium.

### In-cell Photo-affinity labeling (PAL) and click chemistry

Palbociclib-PAL probe was synthesized in a single pot reaction without further purification using an NHS-activated minimalist photo-click compound (CAS 2012552-32-6, Enamine, EN300-28319299). To this end, 10 µl of 100 mM palbociclib in water (CAS 571190-30-2, MedchemExpress, HY-50767) were mixed with 11 µl 100 mM photo-click freshly prepared in water, 1 µl triethylamine added and incubated at 20 °C and 800 rpm shaking for 20 h. Completion of the reaction was analyzed by mass spectrometry, typically confirming >99 % conversion.

Palbociclib-PAL probe was added at 1 µM concentration from a 1 mM stock in DMSO to the media of 70-80 % confluent HeLa cells growing on a 10 cm culture dish and normal cell culture continued for 1 h. Cells were transferred into the UVEN device and immediately irradiated for 0.5-10 s in their original media. For bulb irradiation at a comparable distance, the device was inverted, and culture dishes were propped up on both sides with dedicated placeholders at a distance of 3.5 cm from the UV bulbs, allowing irradiation for 125–10,000 s while the cells remained in their original media. In both cases, cells were washed twice with 10 ml ice-cold PBS, which was removed to completion. Cells were scraped into 500 µl ice-cold, denaturing lysis buffer (25 mM HEPES pH=7.5, 150 mM NaCl, 2 mM MgCl_2_, 0.1 % NP40, 8 M urea) and sonicated three times for 10 s with a Sonopuls ultrasonic homogenizer mini20 (BANDELIN) at 50 % energy on ice with thorough vortexing and spinning down in between. Lysates were centrifuged at 4 °C with 15,000 g for 15 min and supernatants transferred to fresh tubes. The protein content was determined by BCA assay and protein concentrations adjusted to 1 mg/ml with denaturing lysis buffer. Fluorescent dye Cy5.5-azide (Jena Bioscience) was clicked to 50 ug protein lysate by addition of 2 µM Cy5.5-azide, 5 mM CuSO4, 5 mM THPTA, 10 mM sodium ascorbate, and 10 mM aminoguanidine for 2 h at 20 °C and 800 rpm shaking. To remove excess salts for gel analysis samples were acetone precipitated and pellets resolubilized in SDS-PAGE running buffer.

Native lysates were produced from untreated HeLa cells washed with ice-cold PBS, which was removed to completion. Cells were scraped into 200 µl ice-cold, native lysis buffer (25 mM HEPES pH=7.5, 150 mM NaCl, 2 mM MgCl_2_, 0.1 % NP40), sonicated, centrifuged and adjusted to 10 mg/ml with native lysis buffer as described above. Pre-clicked palbociclib-PAL-Cy5.5 conjugate was added to a final concentration of 1 µM concentration in to 25 µg of HeLa native lysate in 10 µl reaction volume and incubated at 20 °C for 30 min with 300 rpm shaking. Tubes were irradiated with UV at a distance of 3.5 cm from the light source (identical distance to LED or bulb). For SDS-PAGE analysis approx. 10 ug of HeLa protein were resolved on a Bis-Tris 4-12 % gradient gel (invitrogen) and imaged on an Odyssey IR scanner (LICOR).

### *In vitro* crosslinking of AIFM1

The gene encoding apoptosis-inducing factor, mitochondrial (AIFM1) with an N-terminal 6×His tag was expressed in Escherichia coli strain BL21(23). Cells were harvested and resuspended in buffer containing 50 mM Tris-HCl (pH 7.8), 500 mM NaCl, 4 mM MgCl₂, 0.5 mM tris(2-carboxyethyl)phosphine (TCEP), 30 mM imidazole, and protease inhibitors (Roche). Cells were lysed by sonication on ice. Debris was removed by centrifugation, and the protein was purified by Ni-NTA affinity chromatography. For further purification, size-exclusion chromatography was performed using a HiLoad 16/600 Superdex 200 pg column (GE Healthcare) equilibrated with 10 mM Tris-HCl (pH 7.8), 150 mM NaCl, and 0.5 mM TCEP. The eluted protein was concentrated to 52 mg/mL, then rebuffered into crosslinking buffer (10 mM HEPES-NaOH, pH 7.8, 150 mM NaCl, 4 mM MgCl₂, 0.5 mM TCEP) and diluted to 1 mg/mL. To induce dimerization, the sample was incubated on ice with 1 mM NADH for 1 h. The crosslinking reaction was performed similarly to previous reports(24). Sulfo-SDA was freshly prepared in crosslinking buffer at 2 mM. Nine separate 23 µg aliquots of AIFM1 (1 mg/mL) were incubated on ice with 0.5 mM Sulfo-SDA for 1 h. To quench the NHS-ester reaction, 50 mM Tris-HCl was added and incubated for 15 min. Each 30 µL sample was pipetted into the centre of a well in a Greiner Bio-One polystyrene 24-well cell culture plate and irradiated with UV light using the UVEN device at maximum intensity for 0, 2, 4, 8, 16, 32, 64, 128, and 180 s. The crosslinked protein samples were then separated by NuPAGE™ 4–12% Bis-Tris gel electrophoresis using MOPS buffer at 180 V. Bands corresponding to monomeric and dimeric AIFM1 (from the 32-second activation sample) were excised and digested in-gel with trypsin, as previously described (25). The resulting peptides were desalted using C18 StageTips(26) prior to LC-MS/MS analysis.

### In-cell crosslinking with L-photo-leucine

The stable HEK293 Expressing Rpn11-HTBH Cell Line (ABM Cat# T6007) that incorporates L-photo-leucine (27) was grown in 145 mm diameter dishes (Greiner Cat# 639160) in DMEM (High glucose, 10% FBS). Cells growing with normal L-leucine instead of L-photo-leucine were used as a control. For harvesting, cells were washed twice with warm PBS and 10 mL ice-cold PBS was added to the plates. L-photo-Leucine activation occurred with UVEN by irradiating twice at maximum intensity for 10 sec, with a 10-sec pause in between. Cells were scraped, centrifuged @200 x g for 5 min and boiled in Laemmli buffer. Samples were run on 4–20% Mini-PROTEAN® TGX™ Precast protein gels (Bio-Rad Cat# 4561096) and blotted on PVDF membranes (Bio-Rad Cat#1704157). After blocking the blot in 5% BSA in TBST, incubating for 30 min with streptavidin-HRP (1:20,000, Thermo Fisher Cat# 21126), and washing 6x with TBST, the biotinylated signals were detected using SuperSignal™ West Pico PLUS Chemiluminescent Substrate (Thermo Fisher Cat# 34580) using a Bio-Rad ChemiDoc XRS+ system.

### Extraction of protein-crosslinked DNA (XDNAX)

The extraction was performed similar to as described recently with slight variation. In brief, cells from one confluent 15 cm diameter culture dish were lysed in 1 ml TRIZOL (T9424, Sigma Aldrich) by pipetting until completely homogenous, combined with 200 µl chloroform, centrifuged with 12,000 g for 10 min at 4 °C. The aqueous phase was discarded and the interphase transferred to a fresh tube to be disintegrated in 1 ml recovery buffer (Tris-Cl pH=7.5 50 mM, EDTA 1 mM, SDS 1 %) until completely dissolved. The DNA was isopropanol precipitated and rehydrated in 900 µl in water with 0.1 % SDS. RNA was digested with 0.5 ug RNase A and T1 for 60 min at 37 °C, 700 rpm shaking. DNA was fragmented by Covaris sonication in 120 µl portions using the parameters 50 cycles, Scan Speed: 1.0, PIP: 300, CPB: 50, AIP: 75, Dithering: Y=1 Speed=10. The sonicated sample was cleared by centrifugation with 5,000 g for 5 min at room temperature. SDS concentration was adjusted to 2 % and the sample incubated at 95 °C for 5 min, 700 rpm shaking. The sample volume was doubled by addition of guanidinium thiocyanate (GuTC) 5 M, and again incubated at 95 °C for 5 min, 700 rpm shaking. The volume was doubled again by addition of ethanol 100 %, and samples applied with 3,000 g to an RNeasy Midi Column (Quiagen). Columns were washed twice with 4 ml buffer RW and three times with buffer RPE. Crosslinked protein-DNA complexes column were eluted overnight with 300 µl nuclease elution mix (NEB nuclease P1 (NEB) buffer 1 x, MgCl_2_ 5 mM, 0.5 µl NEB nuclease P1, 0.5 µg benzonase (SantaCruz)) and again eluted with 300 µl elution buffer (Tris-Cl 50 mM, SDS 2 %). For protein cleanup 10 µl SP3 beads (GE44152105050250, Sigma Aldrich, original slurry) were added and vortexed before addition of 1 ml ethanol 100 %, mixing and protein aggregation for 15 min. The beads were washed four times with 2 ml ethanol 70 % and subsequently spun down to remove all residual ethanol. Protein was digested off the beads overnight in 200 µl trypsin digestion buffer (EPPS 50 mM pH=8, 5 mM DTT, 0.5 µg trypsin per sample) 37 °C, 700 rpm shaking. Cysteines were alkylated by addition of 6 µl chloroacetamide 550 mM while the incubation was continued for 60 min. The beads were collected on a magnet for 5 min and the supernatant transferred to a fresh vial. Peptides were cleaned up by SCX StageTip followed by C18 StageTip into six high-pH fractionations and combined into four fractions with F1 5 + 50 %, F2 10 %, F3 15 %, F4 0+20 % acetonitrile.

### Extraction of protein-crosslinked RNA (XRNAX)

A variation of the previously reported purification was applied. Cells from one confluent 15 cm diameter culture dish were lysed in 1 ml TRIZOL by pipetting until completely homogenous, combined with 200 µl chloroform, centrifuged with 12,000 g for 10 min at 4 °C. The aqueous phase was discarded and the interphase transferred to a fresh tube to be disintegrated in 900 µl recovery buffer (Tris-Cl pH=7.5 50 mM, EDTA 1 mM, SDS 1 %) until completely dissolved. Samples were combined with 100 µl NaCl 5 M and 1 ml isopropanol, mixed and incubated at -20 °C for 30 min. The precipitated sample was spun for 15 min with 17,000 g at -11 °C. The resulting pellet was washed with 2 ml EtOH 100 % and ethanol residues removed to completion. The pellet was then detached from the wall and rehydrated in 430 µl ultrapure water at 4 °C for 15 min on a rotating wheel. The hydrated cloud was further dissolved by pipetting before addition of 50 µl DNase I buffer 10 x (NEB), 1 µl RNaSin and 20 µl DNase (NEB). DNA was digested for 30 min at 37 °C, 700 rpm shaking, remaining clumps dissolved by pipetting and the digestion resumed for another 15 min. Samples were combined with 1.5 ml Quiagen RNeasy lysis buffer (Quiagen) and heated to 80 °C for 10 min at 500 rpm shaking. After they reached room temperature samples were centrifuged with 12,000 g for 5 min and 1900 µl transferred to a fresh tube leaving precipitates behind. They were combined with 1 ml ethanol 100 %, mixed and applied to an RNeasy Midi spin column at 3,000 g for 2 min at room temperature. Columns were washed twice with 4 ml buffer RW and three times with buffer RPE. Protein-crosslinked RNA was then eluted twice with 300 µl ultrapure water. RNA was digested overnight at 37 °C, 700 rpm shaking after addition of 30 µl Tris-Cl 1 M along with 0.5 µg of the RNases A (Sigma Aldrich), T1 (Sigma Aldrich), benzonase (SantCruz) and 5 µl of MgCl_2_ 1 M. For SP3 cleanup of proteins 100 µl of SDS 20 % was added and samples denatured at 90 °C for 5 min. After reaching room temperature 10 µl of SP3 beads were added (GE44152105050250, Sigma Aldrich, original slurry) along with 1 ml ethanol 100 % and protein aggregation allowed to occur for 15 min. Beads were collected on a magnet, washed four times with 2 ml ethanol 70 % and all residual ethanol removed. Proteins were digested off the beads in 200 trypsin digestion buffer (EPPS 50 mM, 5 mM DTT, 0.5 µg trypsin) overnight at 37 °C, 700 rpm shaking. For cysteine alkylation 6 µl CAA 550 mM were added and incubation continued 60 min. Beads were collected on a magnet and 200 µl of the digested peptides mixed with 10 µl formic acid 10 % in a fresh tube. Samples were centrifuged with 20,000 g for 5 min at room temperature, and 200 µl of the acidified peptides transferred to a fresh tube. Peptides were cleaned up by SCX StageTip (225166889-U, Sigma Aldrich), dried down, and desalted by C18 StageTip (66883-U, Sigma Aldrich).

For the benchmarking of this protocol, we selected a set of RNA-binding compounds including translation inhibitors (cycloheximide CHX, harringtonine HAR, puromycin PUR), a series of tetracyclines (tetracyclin TET, doxycycline DOX, minocycline MIN, neocycline NEO), and splicing/ translation modulators (ataluren ATA, risdiplam RIS, branaplam BRA, isoginkgetin ISO). All compounds were purchased from Hycultec, except cycloheximide from Sigma Aldrich. Cells were photolabeled by addition of 100 µM 4SU to the medium for 24 h. All drugs were added to a final concentration of 10 µM concentration from a 1:1,000 stock in DMSO. After five min of incubation at normal culture conditions, cell culture dishes with media were transferred to the UVEN device, irradiated for 5 s, and cells immediately harvested on ice.

### HPLC-MS analysis of proteomic samples

For protein-DNA and protein-RNA crosslinking samples, analysis via data-dependent acquisition (DDA) occurred on an Orbitrap Eclipse Tribrid mass spectrometers (Thermo Scientific), connected to a Dionex UltiMate 3000 RSLCnano system (Thermo Scientific). Samples were injected onto a trap column (75 µm x 2 cm) packed with 5 µm C18 resin (Dr. Maisch Reprosil PUR AQ) in solvent C (formic acid 0.1 %). Peptides were washed with solvent C at 5 µl/min for 10 min and subsequently transferred on to an analytical column (75 µm x 48 cm, heated to 55 °C) packed with 3 µm C18 resin (Dr. Maisch Reprosil PUR AQ) at a flow rate of 300 nl/min using a gradient of solvent solvent from 4 % B (formic acid 0.1 % in acetonitrile, DMSO 5%) followed by a linear increase to 32 % B in A (formic acid 0.1 % in ultrapure water, DMSO 5%). Nanosource voltage was 2,000 V, ion transfer tube temperature 275 °C. Detection occurred with data-dependent acquisition using an OT-OT method and a cycle time of 2 s. MS1 resolution was 60,000, scan range 360-1300, RF lens 40 %, AGC target 100 % and maximum injection time 50 ms. MS2 isolation occurred with a quadrupole window of 1.2 m/z and fragmentation with 30 % HCD energy. MS2 resolution was 30,000, first mass 100 m/z, AGC 200 % and maximum injection time 54 ms.

For protein-DNA and protein-RNA crosslinking samples, analysis via data-independent acquisition (DIA) occurred on a timsTOF HT mass spectrometer (Bruker), connected via a CaptiveSpray source (Bruker) to a Vanquish Neo UHPLC-System (Thermo Scientific). Samples were loaded onto an analytical Aurora Ultimate CSI column (75 µm x 25 cm) packed with 1.7 µm C18 resin (Ionopticks, #AUR3-25075C18-CSI) under pressure control at a maximum of 1,000 bar. Peptide separation was performed at a flow rate of 250 nl/min using a gradient of solvent from 9 % B (formic acid 0.1 % in acetonitrile) followed by a linear increase to 28 % B within 19.8 min, followed by a linear increase to 37% B within 4.5 min in A (formic acid 0.1 % in ultrapure water). An end plate offset of 500 V, capillary voltage of 4500 V and a dry temperature of 180°C were applied. Data were acquired in DIA mode utilizing a 3×8 dia-PASEF window scheme covering a mobility range between 0.64 to 1.45 V*s/cm^2^. A ramp time of 100 ms and advanced collision energy settings were used, resulting in a cycle time estimate of 0.95 s.

Protein-protein crosslinking samples were analyzed on an Orbitrap Astral Mass Spectrometer (Thermo Fisher Scientific) connected to a Vanquish Neo UHPLC System (Thermo Fisher Scientific). Peptides were resuspended in a solution containing 1.6% v/v acetonitrile and 0.1% v/v formic acid and were injected onto a 5.5 cm High Throughput µPAC™ Neo HPLC Column (Thermo Scientific) operating at 50°C column temperature. The mobile phase consisted of water with 0.1% v/v formic acid (mobile phase A) and 80% v/v acetonitrile with 0.1% v/v formic acid (mobile phase B). Peptides were eluted at a flow rate of 300 nL/min using a 50-min gradient: a linear increase of mobile phase B from 2% to 12,5% over 10 min, followed by an increase to 45% in 80 min, then to 55% in 2.5 min, and finally a ramp to 95% B in 2.5 min. Eluted peptides were ionized by an EASY-Spray source (Thermo Scientific) and introduced directly into the mass spectrometer. The mass spectrometry data was acquired using a data-dependent acquisition (DDA) mode with the top-speed option. For each 2.5-second acquisition cycle, the full scan mass spectrum was recorded in the Orbitrap with a resolution of 120,000 resolution with a m/z range of 400 to 1450. The AGC was set to 100% with a maximum injection time of 50 ms. Ions with charge 3-7 and minimum intensity of 1E4 were then individually isolated with a 1.4 m/z isolation window and fragmented using higher-energy collisional dissociation (HCD) with a normalized collision energy of 30%. The fragmentation spectra were then recorded in the Astral mass analyzer. The AGC was set to 100% with a maximum injection time of 20 ms. Dynamic exclusion was enabled with single repeat count and 30 second exclusion duration.

### MS database search

For protein-RNA and protein-DNA crosslinking, MS data acquired in data-dependent (DDA) mode was searched with MaxQuant (28) (2.4.2.0). All searches were performed against the Uniprot human proteome (search term: ‘reviewed:yes AND proteome:up000005640’, 20216 entries, retrieved 13 June 2020). For searches and data processing outside of MaxQuant, its contaminants.fasta was included in the Uniprot human proteome file for consistency with the flag ‘CON_’. MaxQuant settings were kept at their default value, except activation of the iBAQ quantification and setting peptide and protein identification FDR to 1 for PROSIT rescoring (29). For the RNA-interacting proteomes responding to RNA-binding drugs, files were searched in three batches, each processed on the same day, with the ’match between runs’ option enabled. Search results were rescored using an in-house pipeline incorporating PROSIT and Picked Protein Group FDR (30).

For protein-RNA and protein-DNA crosslinking, MS data acquired in data-independent (DIA) mode was searched with DIA-NN (31) (1.9.1). The option ‘FASTA digest for library-free search / library generation’ was selected accepting all automatic default setting. For the analysis of RNA-interacting proteomes extracted after different irradiations (Figure 3B), each file was searched individually to avoid the transfer of identifications. In the case of the RNA-interacting proteomes responding to RNA-binding drugs, files were searched in three batches analogously to the MaxQuant searches and with activated ‘MBR’ to allow for transfer of identifications within one experiment.

For protein-protein crosslinking, MS raw data were converted to peak lists using the MSConvert module in ProteoWizard (version 3.0.11729). Precursor and fragment m/z values were recalibrated. Identification of crosslinked peptides was carried out using xiSEARCH software (https://www.rappsilberlab.org/software/xisearch, version 1,7,6,4) (32). The peak lists were searched against sequences of the expressed AIFM1. The reversed protein sequence of the protein subunits was used as a decoy during the search for error estimation.The following parameters were applied for the database search: MS accuracy, 3 ppm; MS2 accuracy, 5 ppm; enzyme, trypsin (with full tryptic specificity); allowed number of missed cleavages, 2; missing monoisotopic peak, 2. For all samples, carbamidomethylation on cysteine was set as a fixed modification and oxidation on methionine was set as a variable modification. Match to non-covalent links was enabled. The crosslinker was set to SDA. The reaction specificity for SDA was assumed to be for lysine, serine, threonine, tyrosine and protein N-termini on the NHS ester end and any amino acids for the diazirine end. SDA loop link and hydrolyzed SDA on the diazirine end were set as variable modifications. Crosslinked peptide candidates with a minimum of three matched fragment ions (with at least two containing a crosslinked residue) in each crosslinked peptide were filtered using xiFDR (version 2.2.betaB) (33, 34) with the filter setting (scoreP1Coverage >= 3 AND (pepSeq2 is null OR scoreP2Coverage >= 3)) AND (site1>0 OR pepSeq2 isnull) AND ([%fragment unique crosslinked matched conservative%] >= 2.0)”. A false discovery rate of 5% at the residue-pair level was applied. xiVIEW (35) was used to visualize the crosslinking data on the crystal structure of AIFM1 (PDB 4BUR), and to measure the Euclidean distances between Cα atoms of the crosslinked residue pairs.

### Processing of proteomic data

All data analysis was performed in R (4.4) via RStudio (2024.04.2, build 764). For the analysis of proteomic data searched with MaxQuant (DDA acquisition) the proteinGroups_fdr0.01.txt file was used. Contaminants (‘CON_’) and reverse identifications (‘REV_’) were removed. In the case of data searched with DIA-NN (DIA acquisition) the report.pg_matrix.tsv as well as the report.pr_matrix.tsv file were used and contaminants (‘CON_’) removed.

For the comparison of DNA-interacting proteomes between three cell lines, proteins detected with fewer two peptides were removed and iBAQ values median-centered across all replicates. Triplicates of mean iBAQ values were calculated by averaging one of each replicates from each cell line. Missing values were imputed as described before (17) with an adaptation of the function described for PERSEUS (36), using the parameters width 0.3 and downshift 1.8. Negative binomial model-based (NBM) testing occurred with DESeq2 (37) under the assumption that iBAQ represent protein copy numbers on DNA (38). Triplicates of each cell line were tested against triplicates of the mean. Fold changes were corrected using the apeglm package.(39)

To determine half-effective crosslinking times (ET_50_), DIA-NN LFQ values were first median-centered using the intensities of glycoproteins (retrieved from Uniprot via ‘CARBOHYD’), which form a constant background of XRNAX, XDNAX and other TRIZOL-based methodology for proteomics such OOPS(9), that is enriched independent of crosslinking. LFQ values were normalized to the maximum and proteins with relative intensity >0.5 in the unirradiated sample removed as background. Time points before and including the maximum were used for fitting a linear model using log-transformed irradiation times. Slopes of the fit were used to calculate half-effective crosslinking times using the formula ET_50_ = log(2) / slope. For comparisons to the full proteome we used a deep MCF7 total proteome that we recently reported (17).

RNA-interacting proteomes were processed and analysed in four batches, each including triplicates of four drug treatments and a DMSO control. For better consistency each batch included its own triplicates of DMSO-treated control cells, which were used for the differential quantification within one batch. Differential testing occurred for each batch separately using DESeq2 and foldchange correction as described above. In the case of DDA analysis, iBAQ values were used and missing data imputed as described above. Results were filtered for proteins detected with two peptides across all samples after NBM testing (Table S4). In the case of DIA analysis, DIA-NN LFQ values were used and we only allowed proteins detected with more than two peptides, in either drug treatment or DMSO, to enter the differential analysis (Table S3). To this end, we used the report.pr_matrix.tsv to parse all proteins for peptides with intensity, and used the replicate with the lowest peptide count to determine if a protein was included. This strongly enhanced the differential analysis, removing extreme and spurious results.

### Functional annotation of proteins and data visualization

Proteins were annotated via ENSEMBL BioMart accessed via the R package ‘biomaRt’. GO enrichment analysis was performed using the GOrilla web interface. Data was plotted using the R package ‘ggplot2’ and ‘ggbeeswarm’. Renderings of the UVEN device was performed in Fusion360 (Autodesk). Figures were partially generated with Biorender. Figures were composed in Adobe Illustrator 2025 (Adobe).

## Results

### High-intensity UV irradiation accelerates photo-reactions by orders of magnitude

In order to increase protein-DNA crosslinking yields with the lowly photo-reactive nucleotide 4-thiothymidine, we recently reported ‘UV irradiation system for ENhanced photo-activation’ (UVEN) - an LED-based irradiation device for high-intensity UV irradiation of culture cells or other biological specimen.(40) UVEN aims to improve the main shortcomings of a standard bulb-based device, which relate to low intensity of the irradiant and only ice as protection from heat and desiccation (Figure S1A). To increase intensity but limit toxicity of UV light towards living cells, the device uses longwave UV with 365 nm wavelength, avoiding photodamage of protein and nucleic acids absorbing light at 254-280 nm.(17) The employed high-powered UV-LEDs emit more than 2,000 times stronger UV light than conventional bulbs (optical, directly measured at the irradiant). In order to irradiate the area of a cell culture dish, sixteen LEDs are mounted on a large aluminum heat sink that is suspended in the irradiant chamber (Figure S1B and S1C). This chamber is isolated by a quartz glass window from the specimen chamber on top, which physically separates irradiant and biological sample into two chambers that can be cooled independently. This allows for very intense irradiation at controlled temperature (Figure S1D).

During uniform illumination (∼2 W/cm^2^), UVEN is able to accelerate photoreactions homogenously across an entire 15 cm cell culture dish by several orders of magnitude compared to conventional UV bulb irradiation. Figure 1B illustrates this on the example of a kinase inhibitor probe for photo-affinity labeling (PAL). LC-MS quantification demonstrated that UV bulb irradiation required more than 250 s, whereas UVEN irradiation required only 0.5 s for photoconversion of more than 90 % of the initial compound, thus, accelerating the photo-activation of the diazarine more than 500-fold. In order to irradiate smaller specimens such as tissue sections, it is possible to move the LEDs closer to the glass bottom, irradiating small areas (16 times 1 cm^2^) with intensities as high as 10 (±0.1) W/cm^2^, 1400-times more intense than a bulb-based device. We found that mouse tissues exhibited surprisingly high 365 nm light penetration under this configuration. When a 0.5 mm brain tissue section was irradiated at 10 W/cm², a residual light intensity of 1 W/cm² was measured (Figure S1E), which is more than 100 times higher than the surface intensity of an unobstructed 365 nm UV bulb (∼7 mW/cm²). This implies that high-intensity 365 nm light could effectively activate photoreactions throughout brain tissue sections thicker than one millimeter.

To make high-intensity photo-activation available to a wide community of researchers, we established a website that collects open-source construction plans, assembly manuals, software, and future versions of the device at www.uven.org.

### Sub-second protein-drug crosslinking in living cells

We tested high-intensity UV photo-activation on four proteomic applications, starting with in-cell protein-drug crosslinking. A commonly applied chemoproteomic approach to characterize drugs is photo-affinity labeling (PAL), which has been widely explored for drug-target deconvolution and fragment-based screens.(22, 41, 42) For PAL, a parent compound is modified with a photoreactive group and a click handle (Figure 1B). The probe binds target proteins in living cells or lysates, before UV activation covalently crosslinks them in a photoradical reaction, allowing for enrichment under highly denaturing conditions. Despite its utility in living cells, PAL often suffers from low crosslinking yields, even with prolonged irradiation of 10-20 min.(43) We used N-hydroxysuccinimide (NHS) chemistry to functionalize the CDK4/6 kinase inhibitor palbociclib in a single-step reaction without cleanup to a PAL probe (Figure S1F) (43), which we applied to living HeLa cells or native HeLa lysates. After irradiation-induced crosslinking, click chemistry was used to add a fluorescent dye, and photo-crosslinked protein-drug complexes were resolved by SDS-PAGE (Figure S1G). Strikingly, high-intensity UV-LED irradiation of both lysates and living cells led to rapid fluorescent labeling of specific proteins after only 0.5 s of photo-activation, while comparable labeling only occurred after minutes of irradiation with conventional UV bulbs (Figure 1C and 1D). Photoligation in lysates was less specific than in intact cells, where primarily bands corresponding to the palbociclib targets CDK4/6 became apparent. In living cells, the labeling intensity after 1 s of high-intensity UV-LED irradiation was similar to the one achieved after UV bulb irradiation for 500 s, which is close to the commonly applied ∼10 min of irradiation used throughout the current literature (22, 43). The background signal increased markedly after 1,000 s of bulb or 10 s of LED irradiation, indicating that there is a trade-off between crosslinking yield and specificity. Overall, high-intensity UV-LED irradiation accelerated protein-drug photo-crosslinking 500-fold, for the first time enabling PAL in living cells under physiological culture conditions.

### High-yield protein-protein crosslinking for structural proteomics

We next turned towards protein-protein crosslinking. In recent years, the use of photo-activatable diazirine groups in crosslinking MS has gained popularity, as it not only increases crosslinking density(44) but also enhances the contrast and resolution of crosslinking data.(45) We first tested the performance of the UVEN irradiation device on diazirine-based protein photo-crosslinking *in vitro*. The isolated human protein AIFM1 forms a dimer in the presence of NADH.(23) We crosslinked the induced AIFM1 dimer using the photo-crosslinker sulfo-NHS-diazirine (sulfo-SDA). SDS-PAGE analysis revealed that a detectable crosslink-stabilized AIFM1 dimer was yielded with as little as 2 s of activation. The yield of the crosslinked dimer increased with longer UV exposure and plateaued at 32 s (Figure 3A). According to previous studies (46, 47) and the manufacturer’s instructions, achieving an optimal sulfo-SDA activation typically requires about 20-30 min using conventional UV-bulb setups. When subjected to LC-MS/MS, the AIFM1 sample crosslinked with 32 s of UV activation yielded a total of 122 identified crosslinks (at 5% FDR at the linkage level), consistent with expectations for such an experimental setup without any enrichment for crosslinked peptides.(23) Ninety-nine of them could be mapped onto the crystal structure of the AIFM1 dimer (PDB 4BUR), with 86 (87%) having Cα–Cα distances below the calculated crosslinking limit of 30 Å (Figure 3B). This high level of agreement with the crystal structure suggests that the activation did not cause evident structural perturbations.

We next explored the potential of high-intensity LED irradiation for protein crosslinking in intact cells. Cultured HEK293 cells incorporating photo-L-leucine (27) were exposed to UV twice, each for 10 s. The reaction was monitored via western blotting of a biotin-tagged protein. As shown in Figure S1H, after UV exposure, the biotin-tagged protein appeared as a single band in the control sample of cells that had grown on regular L-leucine, whereas higher molecular weight bands emerged in the samples of cells that had grown on photo-L-leucine, indicating successful protein-protein crosslinking. In contrast, the same experiment using a UV bulb setup required 12 min of exposure. Additionally, the UVEN device irradiates from below, allowing crosslinking to proceed in the presence of culture medium instead of PBS. This minimizes cellular stress and enables a more physiologically relevant *in vivo* crosslinking workflow. In summary, UVEN effectively activates diazirine-mediated protein-protein crosslinking both *in vitro* and in intact cells. This significantly enhances the temporal resolution of crosslinking MS experiments, particularly those aiming to monitor dynamic changes in protein structure and interactions over time.

### Photo-crosslinked DNA interactomes quantify genomic regulation across cell types

We next turned to protein-DNA crosslinking, where photo-crosslinking has remained largely unexplored due to various technical challenges. We have recently shown that metabolic DNA labeling with the photo-activatable nucleotide 4-thiothymidine (4ST), combined with high-intensity UV irradiation, strongly enhances the crosslinking of DNA-interacting proteins, enabling comprehensive analysis of the DNA-interacting proteome (Figure 2A)(8). To this end, photo-crosslinked protein-DNA complexes can be purified for LC-MS analysis using DNA as an affinity handle, in a process we termed protein-crosslinked DNA extraction (XDNAX) (Figure S2A).

**Figure 2:**
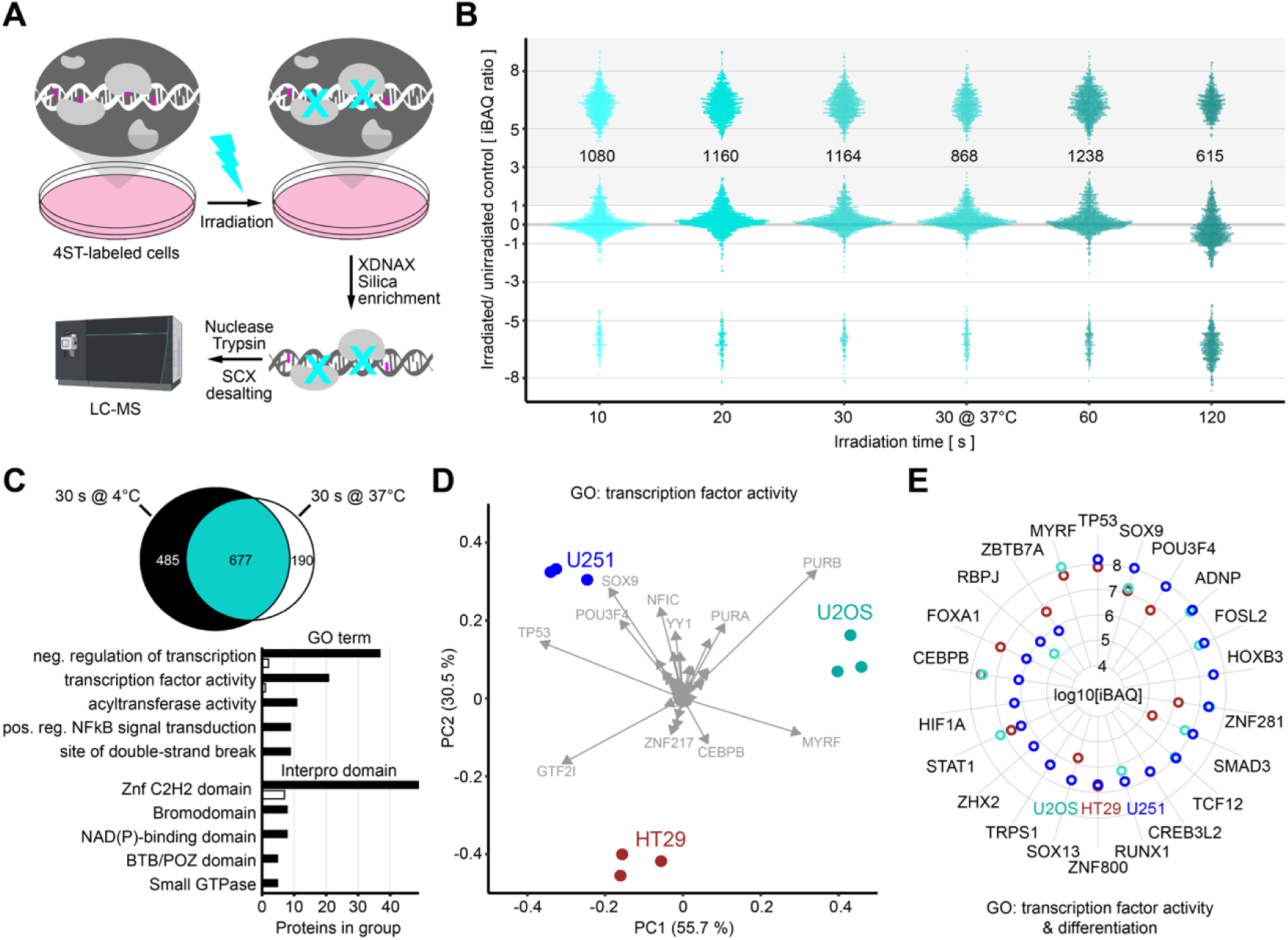
Quantitative comparison of DNA-interacting proteomes from different cell types. A) Workflow for the proteomic analysis of protein-DNA interactions using photo-crosslinking. Cells are metabolically labeled with the photo-activatable nucleotide 4-thiothymidine (4ST), before high-intensity UV irradiation and purification of photo-crosslinked protein-DNA complexes for analysis by LC-MS (XDNAX). B) Bee plot of protein abundances in MCF7 DNA interactomes with increasing UVEN irradiation time relative to unirradiated control cells. All irradiations occurred on cells in ice-cold PBS except for one on cells in their original medium straight from the incubator (30 s at 37 °C). To display proteins without intensity in unirradiated cells pseudocounts were added to iBAQs. C) Venn diagram comparing DNA interactomes of cells UVEN irradiated for 30 s in ice-cold PBS (4 °C) or their original medium (37 °C). Compared are proteins with >10-fold enrichment over unirradiated cells (see B). Barplots show the top 5 GO terms and interpro domains enriched in DNA interactomes derived at 4 °C over 37 °C. D) Biplot for PCA of transcription factor abundances in DNA interactome of three cell lines originating from different germ layers. Replicates of each cell line are shown in same color. E) Radar plot comparing abundances of transcription factors linked to differentiation in DNA interactome from three cell lines.

To better characterize protein-DNA crosslinking kinetics with high-intensity UV, we recorded an irradiation time course in 4ST-photosensitized MCF7 cells. While conventional UV bulb irradiation failed to produce effective protein-DNA crosslinking, high-intensity LED photo-activation as short as 10 s enriched over 1,000 proteins relative to unirradiated controls, 60% of which were Gene Ontology (GO) annotated as nucleic acid-binding (Figure 2B). Extending irradiation beyond 30 s yielded only a marginal increase in DNA-crosslinked proteins (+6.4%), whereas irradiation longer than 60 s caused a marked decrease (−47%), indicating the onset of photodamage. Additionally, 30 s irradiations at 37 °C in cell culture medium resulted in a 36% drop in the number of enriched proteins compared to 4 °C in PBS (Figure 2B), even though the overall abundance of proteins detected under both conditions remained consistent (R² = 0.93, Figure S2B). Comparison of GO annotations for proteins uniquely detected at each temperature revealed a notable enrichment of transcription factors and chromatin regulators, particularly C2H2-type zinc finger proteins, in DNA interactomes crosslinked at 4 °C (Figure 2C), as will be discussed further below.

To evaluate protein-DNA crosslinking across different cell types, we recorded DNA interactomes from three commonly used cell lines derived from different embryonic germ layers (U251 glioblastoma for ectoderm, HT29 colon adenocarcinoma for endoderm, U2OS osteosarcoma for mesoderm), and compared protein abundances in their DNA interactomes by label-free quantification (Table S1). This differential comparison included 1147 proteins with a previous gene ontology (GO) annotation as DNA or RNA binding, 118 of which have been previously reported as human transcription factors.(48) Principle component analysis (PCA) of transcription factor abundances clearly separated the three cell lines and highlighted individual transcription factors contributing most strongly to the separation (Figure 2D). Most variance was explained by GTF2I, MYRF, PURB, SOX9, and TP53, which was undetected on DNA in U2OS cells but among the most abundant transcription factors on DNA in HT29 or U251 cells (Figure S2C and S2D). Notably, both cell lines are homozygous for the dysfunctional TP53 variant Arg273His associated with Li-Fraumeni syndrome, whereas U2OS carries a wildtype form of TP53.(49) The Arg273His mutation occurs at a DNA-contact site within a mutation hotspot of the TP53 DNA-binding domain, and has been reported to change its binding specificity to orchestrate an oncogenic transcriptional program that drives cancer progression.(50) Overall 45 transcription factors showed significant abundance differences between the three DNA interactomes, ten of which were exclusively found in the DNA interactome of U251 glioblastoma cells, including HOXB3, CREB3L2 and HIF1A (p<0.001, negative binomial model (NBM) testing towards mean, Figure S2E). Indeed, independent genome-wide screens in mice and humans have identified HOXB3 and CREB3L2 as drivers of glioblastoma (51, 52), a cancer entity where overactivation of HIF1A often drives severe hypervascularization.(53) We observed very distinct abundances on DNA for transcription factors with involvement in cell differentiation (Figure 2E), and specifically in U2OS cells strongly decreased DNA association of various basal transcription factors involved in RNA polymerase II transcription regulation (Figure S2F). Overall, this comparison illustrates how photo-crosslinked DNA interactomes provide insights into genomic regulation across cell types, revealing quantitative fingerprints of transcription factor activities in direct contact with DNA.

### Rapid photo-crosslinking for the quantification of RNA-interacting proteomes under physiological culture conditions

Finally, we turned to protein-RNA interactions, where various cellular processes, such as transcription(20), splicing(54), ribosome biogenesis(55), or translation(56), occur on a timescale of seconds to minutes. Conventional UV crosslinking with bulbs is therefore typically performed on ice to ‘freeze’ molecular processes during the 1-2 min of irradiation (Figure S1A). We reasoned that faster crosslinking might eliminate the need to ‘freeze’ processes, instead enabling snapshots of protein-RNA interactions under physiological culture conditions, with cells remaining in their original medium.

To test this, we combined metabolic RNA labeling of expanding cells with the photo-activatable ribonucleotide 4-thiouridine (4SU) and high-intensity photo-activation.(5) Moreover, we updated our previously published protocol for the extraction of protein-crosslinked RNA (XRNAX)(8) to analyze photo-crosslinked RNA interactomes via standard label-free proteomics, omitting trypsin predigestion but instead purifying intact protein photo-crosslinked to RNA before standard trypsin digestion (Figure S3A, see methods for details). This simplified the LC-MS analysis by eliminating the need for SILAC, allowing for conventional protein-level quantification by standard DDA and DIA methodology. We compared protein-RNA crosslinking with high-intensity UV-LEDs to conventional UV bulbs, and irradiated 4SU-labeled cells straight from the incubator in their original medium for increasing durations (Figure 3A). Strikingly, a single second of high-intensity LED irradiation led to enrichment of more proteins than 120 s of conventional UV bulb irradiation (Figure 3B). Five s of LED irradiation more than doubled the number of enriched proteins compared to 120 s of UV bulbs, whereas longer irradiation times only marginally improved the outcome. Comparing identical samples analyzed via data-independent acquisition (DIA) with 22 min active gradient time on a timsTOF HT (Figure 3B) to standard data-independent acquisition (DDA) analysis with 60 min active gradient on an Orbitrap Eclipse (Figure S3B), we found that RNA-interacting proteomes were highly amiable to DIA analysis, doubling the number of enriched proteins detected in the crosslinked samples. GO enrichment analysis indicated that this particularly included lower abundant RNA-binding proteins involved in chromatin organization and mitochondrial translation (Table S2). In summary, high-intensity LED irradiation dramatically accelerated protein-RNA crosslinking to just a few seconds allowing for their fixation under physiological culture conditions. In combination with high-sensitivity DIA analysis this enabled quantification of near-complete RNA-interacting proteomes within 30 min analysis time.

**Figure 3:**
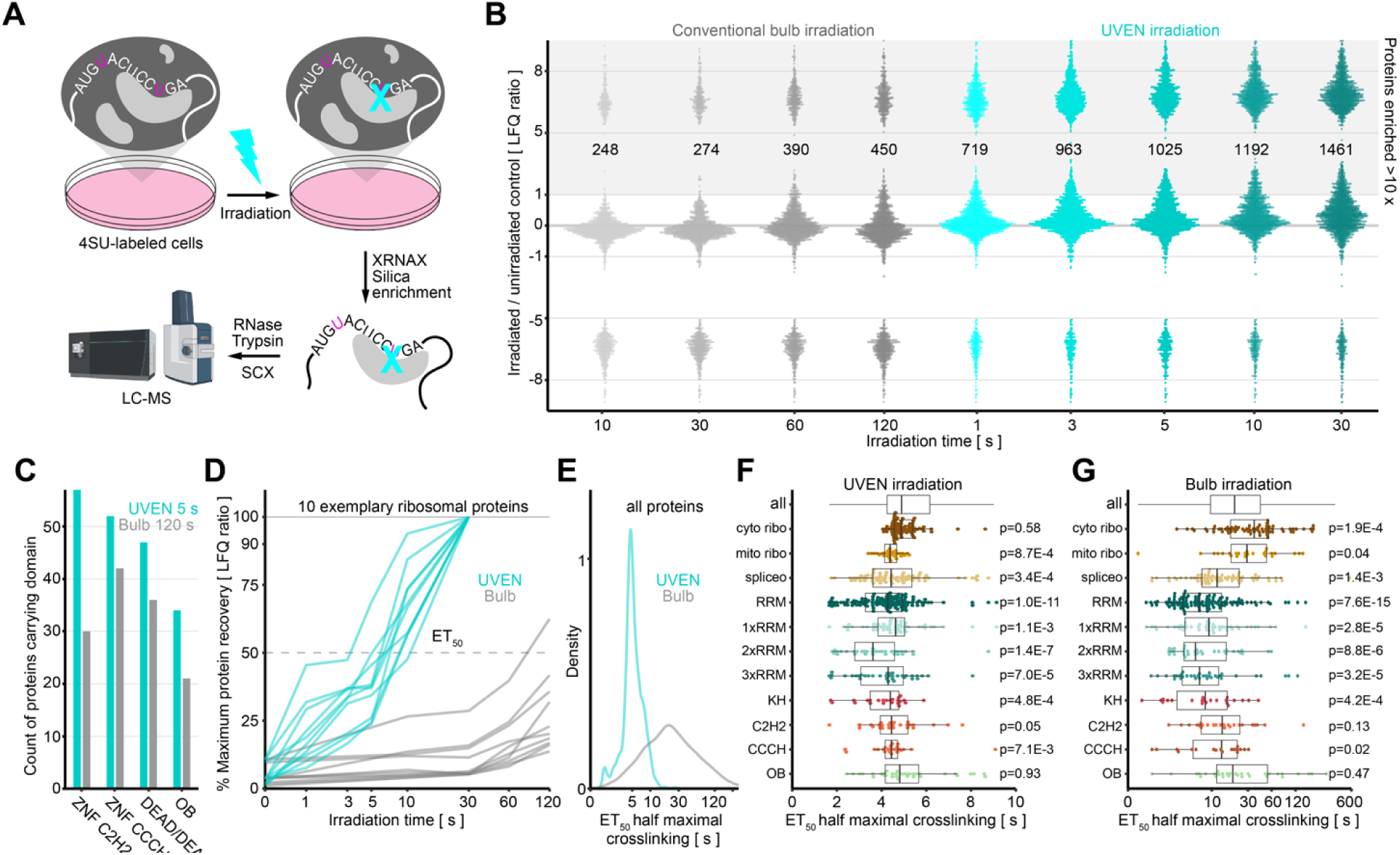
Quantitative analysis of RNA-interacting proteomes and their photo-crosslinking kinetic. A) Workflow for the proteomic analysis of protein-RNA interactions using photo-crosslinking. Cells are metabolically labeled with the photo-activatable nucleotide 4-thiouridine (4SU), before high-intensity UV irradiation in the original medium and purification of photo-crosslinked protein-RNA complexes for analysis by LC-MS (XRNAX). B) Bee plot of protein abundances in RNA-interacting proteomes extracted after increasing irradiation time with a conventional bulb-based device or UVEN. An equivalent of 10 million MCF7 cells was analysed by DIA on a timsTOF HT using a 22-min gradient, see Figure S3B for DDA comparison. To display proteins without intensity in unirradiated cells pseudocounts were added to LFQ values. C) Barplot comparing occurrences of proteins with specific DNA-binding domains between RNA interactomes derived with 120 s bulb or 5 s UVEN irradiation. D) Line plot comparing the recovery of 10 exemplary cytosolic ribosomal proteins after increasing crosslinking time and XRNAX. Each line represents one protein. Normalization occurred towards the maximum LFQ for each protein. E) Density plots for half-maximal crosslinking times under UVEN or conventional bulb irradiation. Compared are all proteins (left panel) or proteins annotated in KEGG as part of the cytosolic ribosome, mitoribosome, spliceosome (right panels). F) Boxplots showing half-maximal crosslinking times between different protein groups under UVEN irradiation. Displayed are all proteins annotated in KEGG as part of the cytosolic ribosome (cyto ribo), mitoribosome (mito ribo), spliceosome (spliceo), proteins carrying RRM domains (RRM), of those carrying one, two or three RRM domains (1,2,3xRRM), K homology (KH), C2H2-type zinc-finger, OB-fold (OB) domains. Testing occurred between all recovered proteins (all) and each group using a Wilcoxon ranksum test and Bonferroni-Holm correction. Note linear scaling. G) Same as in F but for conventional bulb irradiation. Note logarithmic scaling.

### Quantitative and qualitative differences between rapid and slow protein-RNA photo-crosslinking

Next, we asked whether there were differences in the RNA-interacting proteomes derived by conventional bulb or high-intensity LED photo-activation. The absolute protein yields recovered by XRNAX were very similar for common proteins detected both after 120 s of bulb irradiation or 5 s of LED irradiation (Figure S3C and Figure S3D). However, LED-irradiated samples were enriched with many additional proteins not detected at all in bulb-irradiated samples (Figure S3E). This suggested a qualitative advantage in crosslinking with high-intensity UV irradiation. Proteins uniquely enriched after 5 s of high-intensity irradiation were markedly lower in abundance within the RNA-interacting proteome compared to those also identified after 120 s of bulb irradiation (Figure S3F). We compared the abundance distributions of both groups in the MCF7 total proteome and found them to be very similar (Figure S3G), suggesting that the additional crosslinking was unlikely related to protein concentration in the cell. Interaction network analysis via STRING verified that proteins only identified by UVEN were strongly enriched in RNA-binding proteins (FDR=1.1E-78), with the largest interaction clusters relating to mitochondrial translation, translation initiation and mRNA splicing. The most common domain among them was the C2H2-type zinc finger domain, which occurred only half as often after UV bulb irradiation (Figure 3C). Notably, under high-intensity LED irradiation C2H2-type zinc-finger proteins were among the proteins with the best recovery of all proteins in the RNA-interacting proteome relative to the total proteome (Figure S3H).

To assess the crosslinking behavior of all proteins enriched by XRNAX, we compared absolute yields of purified proteins across different irradiation time points using label-free quantification. In most cases we observed logistic growth for the amount of RNA crosslinked protein with increasing irradiation time, allowing us to determine half-effective crosslinking times (ET_50_, Table S2, see Methods for details) (Figure 3D). As expected, ET_50_ values were on average substantially shorter under LED irradiation and spanned a much narrower time window (Figure 3E). This was particularly prominent for ribosomal proteins, which displayed a broad range of ET_50_ values under bulb irradiation but a remarkably narrow distribution under UVEN irradiation (ET_50_ standard deviation σ = 0.7 s vs. 135.7 s; Figure 3F and 3G). We used ET_50_ values to compare different RNA-binding domains and found particularly short crosslinking times for proteins containing RRM (adjusted p = 1E-10, Wilcoxon rank-sum test with Bonferroni-Holm correction) or KH domains (adjusted p = 9.4E-4). Proteins carrying two RRM domains showed very rapid crosslinking (adjusted p = 7.1E-7), the fastest being the poly-U binders CPEB2 and CPEB4, which also exhibited the strongest relative recovery compared to the total proteome (Figure S3H). Importantly, we observed very low correlation between protein abundances in the MCF7 total proteome and ET_50_ values for both bulb (Pearson R² = 0.05) and LED (R² = 0.09) irradiation (Figure S3I). Furthermore, the abundance distributions in the MCF7 total proteome were virtually identical for proteins with short and long ET_50_ values (Figure S3J), again verifying that 4SU crosslinking occurred independent of cellular protein concentration. Overall, these findings showed that rapid protein-RNA crosslinking with high-intensity irradiation provided qualitative and quantitative advantages over slow crosslinking with low-intensity irradiation, which failed for many proteins to produce sufficient yield at physiological temperature.

### Differential quantification of RNA-interacting proteomes upon acute drug action

We benchmarked rapid protein-RNA crosslinking with high-intensity photo-activation on the quantitative analysis of RNA-interacting proteomes from MCF7 cells responding to a panel of twelve RNA-binding drugs. Since only seconds of irradiation were required and cells could remain in their original media during photo-crosslinking, we chose a short treatment time of 5 min to capture immediate changes in protein-RNA interactions during early drug engagement. For LC-MS analysis, we compared two single-shot approaches using the 22-min DIA method and the 60-min DDA method described above. DIA analysis enabled in median differential quantification of 1728 proteins, and DDA 409 (Figure S4A, Table S3 and Table S4, adj. p<0.01, NBM testing, see methods for details). On average, 20 proteins identified by DDA showed significantly altered RNA interactions, compared to 47 proteins with DIA, primarily increasing the quantification of lower abundant proteins (Figure S4B). With a Pearson R^2^ = 0.93, we observed good correlation in fold changes of RNA interaction for proteins identified as significantly regulated by both acquisition methods (adj. p<0.01 in DIA and DDA), in line with a previous report that used conventional UV irradiation and a different extraction method (TRAPP).(57) RNA-interacting proteomes from cells treated with risdiplam showed one of the largest benefits from DIA analysis (Figure S4C), revealing differential RNA interaction for more than double the number of proteins than DDA. This included various proteins involved in RNA splicing, demonstrating that DIA provided additional, functionally relevant information on the activity of the splicing modulator (Figure S4C and S4D). As our drug panel contained compounds known to affect translation and splicing, we focused our analysis on protein constituents of the ribosome and spliceosome. Figure 4A summarizes all drug-induced changes in RNA interactions among constituents of the cytosolic ribosome, highlighting harringtonine, which strongly reduced RNA interactions for a selected set of small ribosomal subunit proteins. Particularly decreased interaction was observed for RPS3 (adj. p=1.1E-4), whose contact with the mRNA at the entry channel is known to stabilize preinitiation complexes at the start codon.(58) This aligned with the known mode-of-action of harringtonine, which traps 80S ribosomes after the formation of the initial peptide bond, blocking the formation of consecutive preinitiation complexes.(56, 59) Turning to the spliceosome, we observed extensive remodeling of protein-RNA interactions for risdiplam and ataluren, as well as somewhat weaker responses for branaplan, minocycline and doxycycline (Figure 4B). Risdiplam and ataluren in particular elicited a strong effect on SR proteins, nine out of 13 of which significantly decreased their RNA interaction under risdiplam (adj. p<0.01, Figure 4C). A direct comparison of the two compounds indicated that both interfered in remarkably similar ways with proteins involved in exon definition and early splice site selection (SNRPA, SR proteins; Figure 4B), while increasing RNA interactions of branch point interactors (U2AF2, PUF60, SF3B1, SF3B3), as well as helicases involved in later splicing stages (DHX16, DHX15, AQR).(60) We speculate that both compounds stabilized structures within the pre-mRNA substrate that required extensive helicase activity for splicing to continue, slowing down steps following 5’ splice site definition. A specific effect of risdiplam was strongly decreased RNA interaction of RNPS1 (adj. p=0.002, Figure S4E), which has been shown to promote exon skipping (61, 62), potentially relating to the ability of risdiplam to promote exon inclusion. Overall, this confirmed that our optimized XRNAX workflow, combined with enhanced photo-crosslinking using the UVEN device, enabled the quantitative analysis of RNA-interacting proteomes with a temporal resolution of minutes under physiological culture conditions.

**Figure 4:**
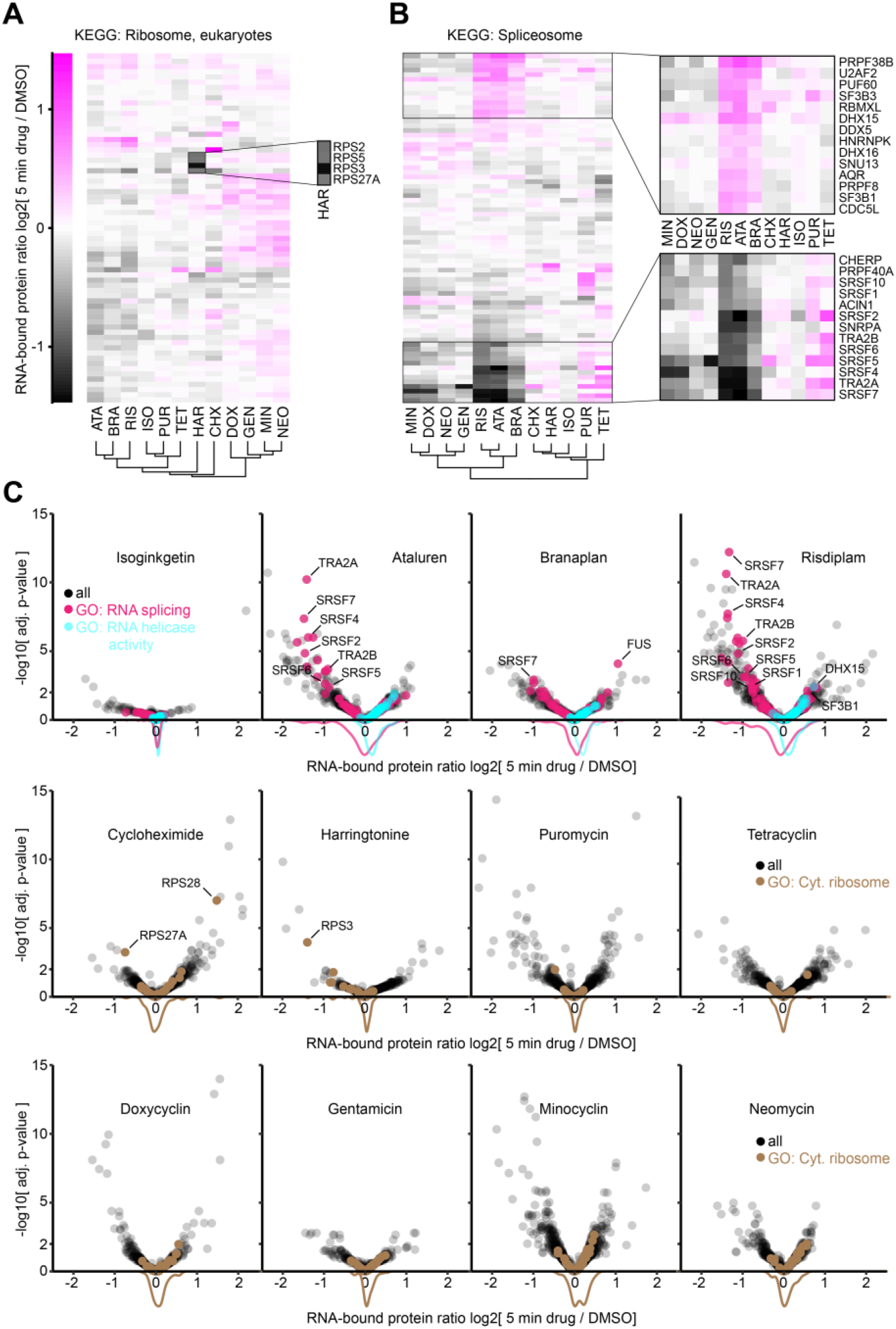
Quantitative comparison of RNA-interacting proteomes in response to RNA-active drugs. A) & B) Heatmaps comparing changes in RNA-interacting proteomes of MCF7 cells treated with 12 RNA-active drugs for 5 min compare to a mock-treated control (DMSO). ATA: ataluren, BRA, branaplan, RIS: risdiplam, ISO: isginkgetin, PUR: puromycin, TET: tetracyclin, HAR: harringtonine, CHX: cycloheximide, DOX: doxycycline, GEN: gentamicin, MIN: minocycline, NEO: neomycin. A) Displayed are all detected constituents of the cytosolic ribosome. Inset magnifies proteins with strongest loss in RNA-interaction. B) Same as in A but for constituents of the spliceosome. Insets magnify proteins with particular strong gain or loss in RNA interaction. B) Volcano plots comparing the effect of 12 RNA-active drugs compared to a mock-treated control (DMSO) for all detected proteins in the RNA-interacting proteome. Density plots indicate fold change distributions of proteins involved in RNA splicing (magenta), RNA helicases (cyan), or the cytosolic ribosome (brown).

## Discussion

### Technical considerations on high-intensity UV irradiation of biological specimens

Conventional bulb-based devices for the activation of photoreactions are limited by their photon output and their ability to protect biological samples during irradiation. One reported setup increased UV intensity by densely packing standard UV bulbs to illuminate an irradiation chamber from both the top and bottom.(63) This approach achieved an intensity of 0.2 W/cm² - approximately a 50-fold increase compared to conventional devices - likely representing the maximal achievable intensity with back-to-back packed UV bulbs without mirrors or lenses. Although lasers in the UVA range with very high intensities exist, their irradiation field is typically no larger than 1 mm, making it prohibitively costly to build an array capable of illuminating a cell culture dish with full intensity.(1, 64–66) Yet, individual UV-lasers and UV-LEDs have been used for photo-crosslinking in small vials.(1, 66–68) UVEN uses an array of UV-LEDs to irradiate an area of over 175 cm² with intensities of around 2 W/cm² - approximately a 500-fold increase compared to conventional bulb devices - without reaching the maximal LED packing density. Further acceleration of crosslinking reactions with more intense UV light could be desirable for in-cell kinetic studies.(1, 66) Employing additional LEDs, future versions of UVEN will increase the intensity by another order of magnitude, potentially pushing most photoreactions in cells to completion in fractions of a second.

### High-intensity photo-activation solves old problems and opens new avenues

We demonstrate that UVEN enhances photo-activation across various disciplines of photo-crosslinking, both *in vitro* and in living cells. UVEN accelerated in-cell photo-activation of a PAL probe from 10 min to just 1 second under physiological culture conditions. This not only implies a substantial gain in throughput but also improves sample handling in a way that paves the way for previously inconceivable PAL-based drug and fragment screens.(22, 42, 43, 69) Similarly, high-intensity photo-activation reduced the time required for effective protein-RNA crosslinking to few seconds while cells remained in their original media. It has been reported that changes in temperature or extracellular osmolality are quickly countered by the cell via formation of molecular condensates that free up intracellular water, thereby avoiding cellular damage or dysfunction.(70) Hence, rapid crosslinking at physiological culture temperature at constant osmolality might prevent spurious interactions that only occur when cells are washed with cold saline or placed on ice.(71) Indeed, using rapid, high-intensity crosslinking, we observed highly consistent quantification of RNA interactomes from cells treated with RNA-binding drugs, demonstrating that minimizing perturbations during the crosslinking process significantly enhances the quantitative analysis of protein-RNA interactions.

### Links between temperature and crosslinking efficiency of proteins with nucleic acids

During the analysis of DNA and RNA-interacting proteomes, we observed that for a group of proteins crosslinking efficiency was sensitive either to temperature or UV intensity, often affecting zinc-finger proteins of the C2H2-type (Figure S4F). Proteins showing sensitivity in both DNA and RNA crosslinking contained between five and 15 C2H2-type domains per protein, suggesting that impaired crosslinking was not due to a low number of C2H2-type repeats. Recent studies using nucleotide-peptide hybrids to map DNA(17) or RNA(72, 73) interactions at site-level resolution reported very few crosslinks for zinc-finger proteins in their C2H2-type domains but primarily in flexible linkers between C2H2-type repeats or other intrinsically disordered regions. Interestingly, it has been reported that the flexible linkers between C2H2-type domains are critical for DNA binding and their phosphorylation is a common mechanism for deactivation of C2H2-type zinc-finger proteins during mitosis.(74) Intrinsically disordered sequences typically interact with nucleic acids in a variety of conformational states (75), only some of which can lead to a successful crosslink. We speculate that in the case of C2H2-type zinc-finger proteins, this results in conformational gating of the crosslinking reaction at physiological temperature, when molecular motion is high and conformations change rapidly. Nevertheless, crosslinking can still be achieved by using a sufficiently reactive photoprobe - such as 4SU, but not 4ST - in combination with high photon flux during irradiation - as provided by high-intensity UV-LED but not by conventional UV bulbs. This illustrates how crosslinking of unstructured regions with low nucleic acid affinity can be affected by nearby high-affinity nucleic-acid binding domains, temperature, and light intensity.

## Data Availability

Proteomic data and search results have been deposited in the MassIVE database under the identifier MSV000097903.

## Author Contributions

Conceptualization, J.T. and B.K. methodology, J.T., P.P., S.T., L.H., Z.C.; investigation, J.T., P.P., S.T., L.H., Z.C., M.M., M.S.; writing—original draft, J.T., B.K., J.R., Z.C.; resources, B.K, J.R.; data curation, J.T.; writing—review and editing, all authors; visualization, J.T., Z.C.; supervision, B.K., J.R.; project administration, J.T.; funding acquisition, J.T., J.R., B.K..

## Acknowledgements

We thank all members of the Kuster laboratory for continuous support and discussion, in particular Karl Kramer, Johanna Tüshaus and Guillaume Medard. We thank Johanna Tüshaus, Xintong Sui and Julian Müller for help with the MS measurement. We thank Patroklos Samaras and Tobias Schmidt for help with the initial LED crosslinking tests. We thank Nicholas Kuhn (UCSF) and Christian Frese (Bayer) for advice and discussion. We gratefully acknowledge funding by the German Research Council DFG supporting this work (DFG project number 492625837). We also acknowledge funding from the Bavarian research foundation for construction of the UVEN irradiation system (BFS project number AZ-1444-20C).

---

**Figure S1:**
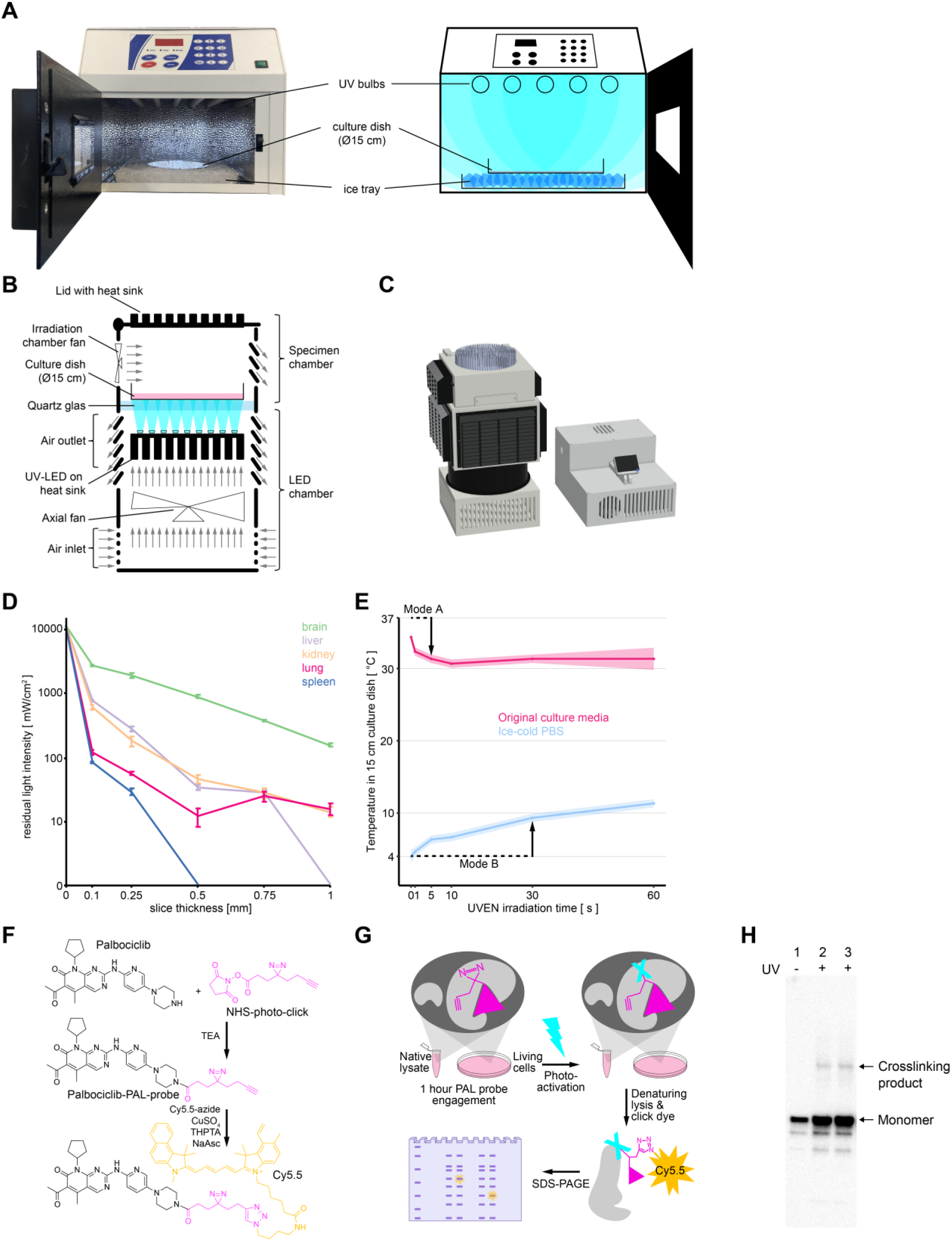
UV irradiation devices for the activation of photo-reactions in biological specimens. A) Photo and schematic of a standard, bulb-based UV irradiation device. To irradiate adherent cultured cells, the medium is removed, and the culture dish is placed on an ice tray positioned beneath an array of UV bulbs. B) Schematic representation of the UVEN irradiation device. High intensity UV-LEDs serve as irradiant, which illuminate a biological specimens through a glass window from below. C) Rendering of the UVEN irradiation device and its control unit. The power supply and control electronics are located in a second housing (right) separate from the irradiation tower (left, see also B). D) Line plot displaying the temperature development of 20 ml culture medium (red, 37 °C initially) or PBS (blue, 4 °C initially) in a 15 cm diameter culture dish during UVEN irradiation. E) Line plot showing UV transmission from a high-intensity UV-LED through mouse tissue slices of increasing thickness, mounted between two glass slides. A light-tight blind was used to eliminate stray light, ensuring that only light passing through the tissue slice reached the UV meter (LS-128, Linshang). F) Chemical structures for the reaction of the kinase inhibitor palbociclib to a PAL probe, and further derivatization with a fluorescent dye (Cy5.5) via copper-catalyzed click chemistry for in-gel imaging. G) Fluorescence imaging of SDS-PAGE after PAL in native lysates. Compared are various time points of LED irradiation (UVEN) to conventional bulb irradiation at identical distance. H) Western blot analysis of L-photo-leucine–mediated crosslinking in intact cells. The biotin-tagged protein appears as a single band under the control condition without L-photo-leucine incorporation (lane 1). Higher molecular weight crosslinked products are observed in the L-photo-leucine–incorporated samples (two replicates, lanes 2 and 3) upon 20 s of UV activation.

**Figure S2:**
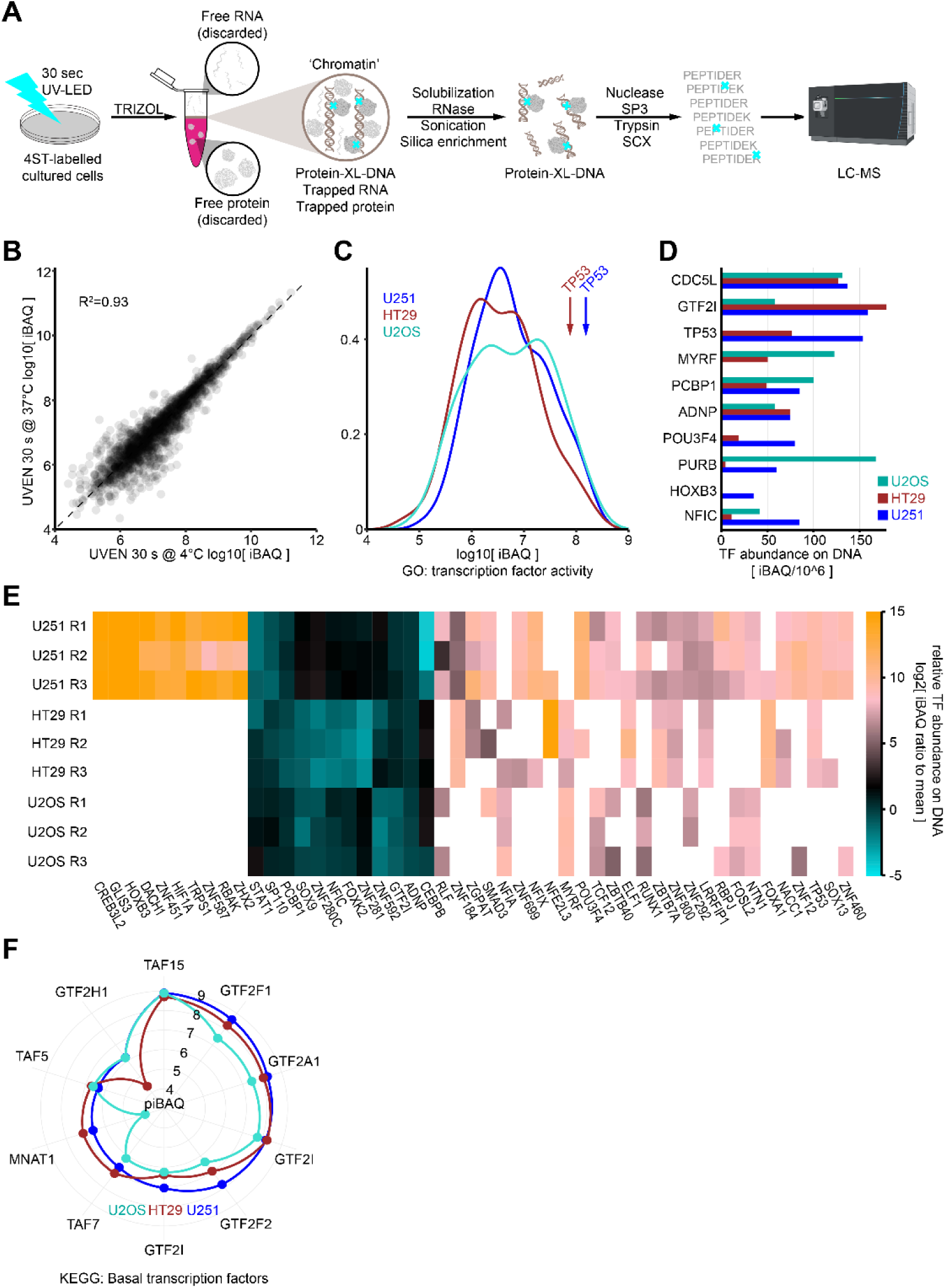
Comparison of transcription factor activity via DNA-crosslinked proteomes from three cell lines. A) Schematic for the extraction of protein-crosslinked DNA (XDNAX) via TRIZOL extraction and silica column purification. B) Scatter plot comparing the abundances of proteins in DNA-interacting proteomes derived from cells crosslinked at 4 °C versus 37 °C. C) Density plot showing the distribution of transcription factor abundances in DNA-interacting proteomes from three different cell lines. Mutant TP53 is abundantly detected on DNA in HT29 and U251 cells, while wild-type TP53 in U2OS cells remains undetected. D) Bar plot comparing the abundances of ten transcription factors showing the most significant differences across the three cell lines. A negative binomial statistical model (NBM testing) was applied to each cell line individually, using the mean across all three as a reference (see Methods for details). E) Heatmap illustrating the abundances of transcription factors with the most significant deviations from the mean (p < 0.001, NBM testing). Protein abundance in each triplicate was normalized to the mean abundance across all cell lines. F) Radar plot comparing the abundances of basal transcription factors in the DNA-interacting proteomes of three cell lines.

**Figure S3:**
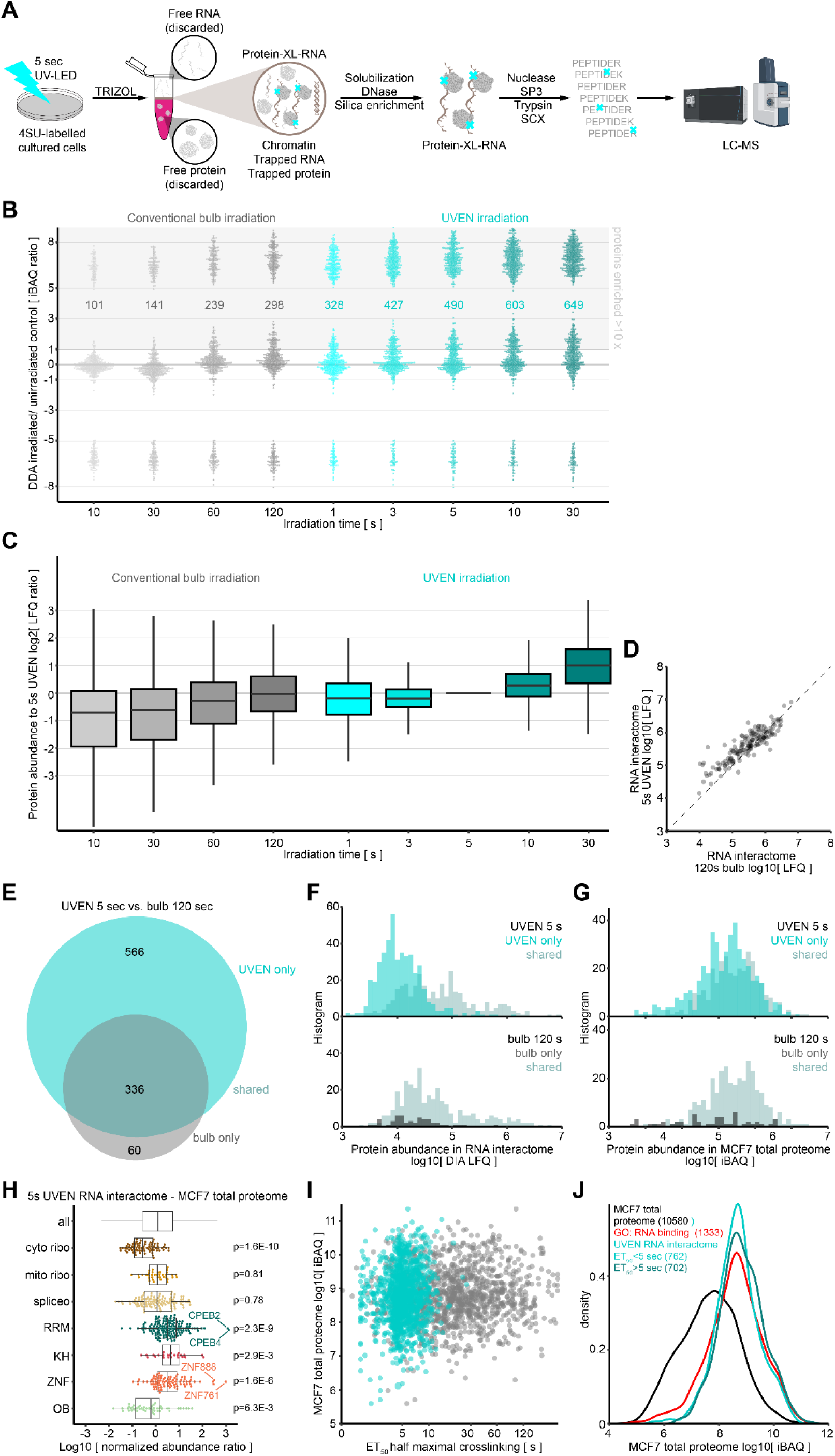
Crosslinking kinetics of protein-RNA interactions using 4SU with low or high-intensity UV. A) Schematic for the extraction of protein-crosslinked RNA (XRNAX) via TRIZOL extraction and silica column purification. Presented is an updated protocol (see Methods for details), that forgoes trypsin predigestion before the silica enrichment to allow for consistent label-free quantification of RNA-interactomes between conditions. B) Beeplot of protein abundances in RNA-interacting proteomes extracted after increasing irradiation time with a conventional bulb-based device or the UVEN. An equivalent of 10 million MCF7 cells was analysed by DDA on an Orbitrap Eclipse using a 60-min gradient, see Figure 3B for DIA comparison. To display proteins without intensity in unirradiated cells pseudocounts were added to LFQ values. C) Boxplots comparing protein abundances between RNA-interacting proteomes normalized to 5 s of UVEN irradiation. D) Scatter plot showing protein abundances of proteins in RNA-interactomes derived after 120 s bulb or 5 sUVEN irradiation (see Figure 3B). Compared are only proteins enriched more than tenfold in both samples compared to an unirradiated control. E) Venn diagram showing overlap between proteins enriched more than tenfold compared to an unirradiated control (see Figure 3B). F) Histogram comparing protein abundances in RNA-interacting proteomes between groups in E. G) Histogram comparing protein abundances in MCF7 total proteome between groups in E. H) Beeplot showing recovery of proteins in RNA-interacting proteomes crosslinked for 5 s with UVEN relative to the MCF7 total proteome. Shown is the ratio of z-scored protein abundances, testing occurred with a Wilcoxon ranksum test between the indicated groups and all proteins. I) Scatter plot comparing half-maximal crosslinking times (ET_50_) with protein abundances (iBAQ) in a deep MCF7 total proteome. Each point represents one protein; UVEN data is shown in cyan, bulb in grey. J) Density plot showing protein abundances in a deep MCF7 total proteome. Compared are proteins with long or short UVEN ET_50_ values, as well as all proteins with a GO annotation for RNA binding.

**Figure S4:**
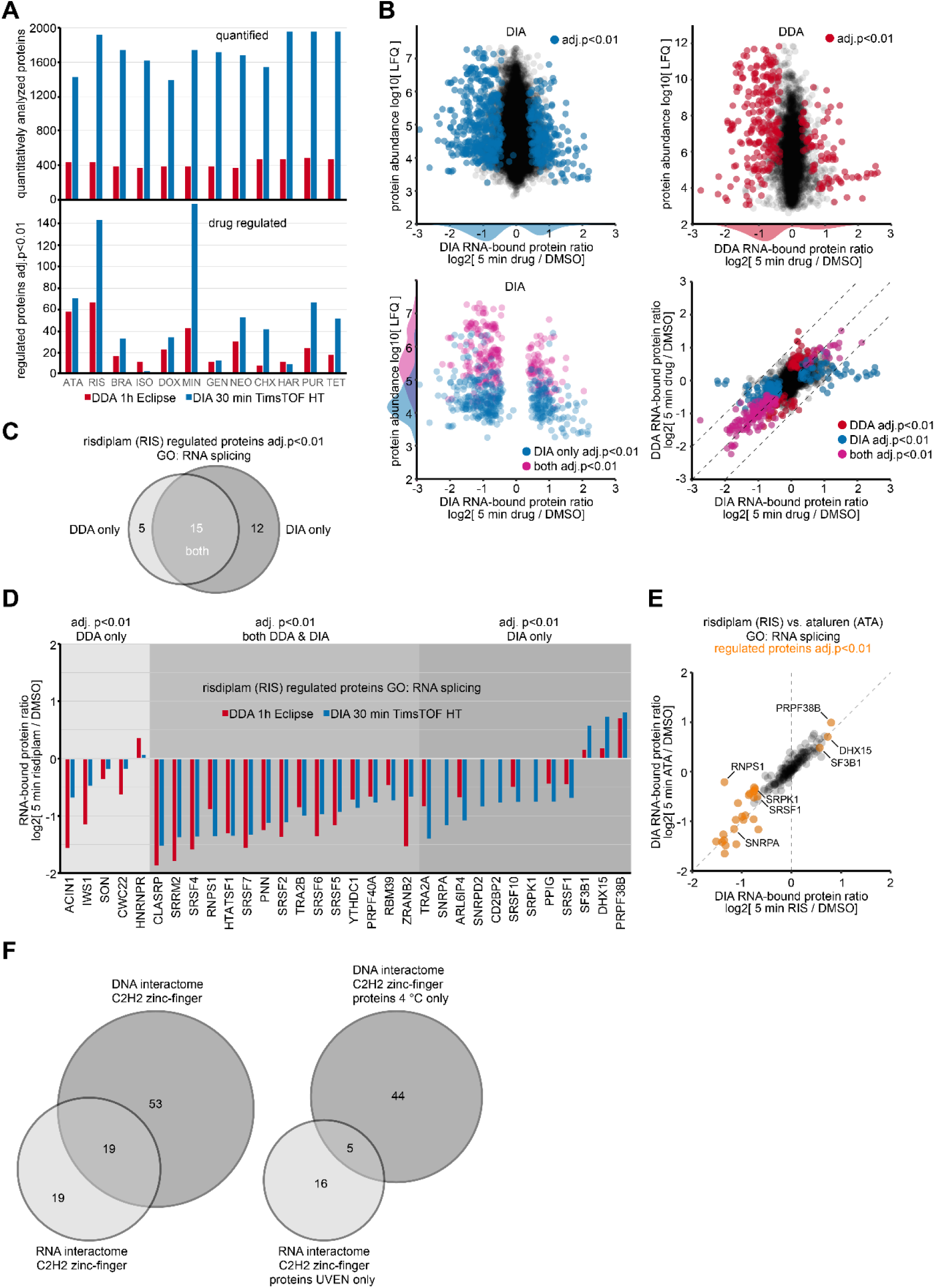
Differential analysis of RNA-interacting proteomes via DDA and DIA methodology. A) Barplots displaying proteins quantitatively compared in RNA-interacting proteomes from cells treated with different RNA-binding drugs. Identical samples were injected on different LC-MS systems for comparison, see Methods for details. Upper panel: All proteins in differential analysis towards mock-treated cells after imputation of missing values for triplicates. Lower panel: Proteins found with significant differences (adj. p<0.01 after NBM testing). B) Comparison of proteins abundances of in RNA-interacting proteomes detected via DDA or DIA analysis. Top left: Scatterplot comparing DIA protein abundances to foldchanges for all 12 drugs combined. Lower left: Same as above only showing proteins identified as significant in DIA analysis (blue), or by both DIA and DDA (purple). Upper right: Scatterplot comparing DDA protein abundances to foldchanges for all 12 drugs combined. Lower right: Scatterplot comparing foldchanges between DIA and DDA differential analysis for all 12 drugs combined. C) Venn diagram of proteins showing significant changes in RNA interaction under risdiplam treatment (adj. p<0.01) with an annotated involvement in RNA splicing. D) Barplots comparing changes in RNA interaction for protein groups from C. E) Scatterplot comparing foldchanges in protein-RNA interactions between risdiplam and ataluren treatment. Only proteins with an annotated involvement in RNA splicing are displayed, proteins with significant change under either treatment highlighted in yellow (adj. p<0.01, NBM testing). F) Venn diagrams showing overlap of C2H2-type zinc-finger proteins between DNA and RNA-interacting proteomes. Left: All C2H2-type zinc-finger proteins derived under optimal irradiation conditions (RNA: 5 second UVEN 37 °C, DNA: 30 second UVEN 4 °C). Right: All C2H2-type zinc-finger proteins exclusively derived under optimal irradiation conditions (not present upon RNA: 5 second UV-bulb 37 °C, DNA: 30 second UVEN 37 °C).

